# TMS-Evoked Corticospinal Beta Oscillations in Humans Recorded from Muscles

**DOI:** 10.64898/2026.02.28.708728

**Authors:** María Sarasquete, Alejandro Pascual-Valdunciel, Federica Ciurluini, Jack de Havas, Sven Bestmann, Dario Farina, Lorenzo Rocchi, Ricci Hannah, Jaime Ibanez

**Affiliations:** BSICoS Group, I3A and IIS, Universidad de Zaragoza, Zaragoza, Spain; Department of Bioengineering, Imperial College, London, UK; Department of Medical Sciences and Public Health, University of Cagliari, Cagliari, Italy; Department of Clinical and Movement Neurosciences, Institute of Neurology, University College London; Department of Imaging Neuroscience, Institute of Neurology, University College London, UK; Centre for Clinical Neuroscience, Hospital Los Madroños, Brunete, Spain; Centre for Human & Applied Physiological Sciences, King’s College London, London, UK; Centro de Investigación Biomédica en Red en Bioingeniería, Biomateriales y Nanomedicina, Instituto de Salud Carlos III, Madrid, Spain

**Keywords:** transcranial magnetic stimulation, corticospinal tract, motor cortex, electromyography, beta rhythm, motor neuron

## Abstract

**Background:** Transcranial magnetic stimulation (TMS) can entrain oscillatory brain activity, offering a promising approach to study motor-related neural oscillations such as beta rhythms. However, how TMS-induced corticospinal oscillations are generated, propagated, and related to endogenous activity remains unclear, partly due to limitations in brain recording techniques. Recording muscle activity provides an alternative and physiologically grounded window into corticospinal dynamics.

**Methods:** We investigated whether subthreshold TMS over the motor cortex induces corticospinal oscillatory activity detectable in muscles, and whether these responses share neural generators with endogenous beta rhythms. Single-pulse subthreshold TMS was applied over the motor cortex in healthy participants while electromyography (EMG) was recorded from tonically active muscles. Stimulation intensity, coil orientation, and stimulation site were systematically varied. Concurrent electroencephalography (EEG) was used to assess cortical responses and corticomuscular transmission. In additional experiments, advanced EMG techniques were employed to track motor neuron pools and characterize how TMS-evoked oscillations are transmitted at the motor unit level.

**Results:** TMS elicited a robust and short-latency increase in beta-band activity in the EMG. The analysis of the elicited muscle responses and the comparison of results across different TMS configurations indicate that the beta responses resulted from activation of inhibitory interneurons in the targeted primary motor cortex. Importantly, the characteristics of cortico-muscular coherence and beta projection to the muscles indicate that the elicited beta responses with TMS have same cortical sources as endogenously generated beta activity.

**Conclusions:** These findings demonstrate that muscle recordings provide a sensitive and physiologically meaningful readout of TMS-induced corticospinal beta oscillations.

## Introduction

Oscillatory neural activity, reflecting the coordinated timing of neuronal population activity, is thought to be a fundamental feature of communication and information processing across distributed networks of the central nervous system [1]. In the motor system, beta-band rhythms represent a prominent signature of motor corticospinal and cortico-basal ganglia activity [2–6] that is thought to play a critical role in movement control. A major obstacle to advancing our understanding of these rhythms and their role in controlling muscle activity is that they are typically intermittent and not tightly time-locked to external events [7,8].

Transcranial magnetic brain stimulation (TMS) offers a means to experimentally evoke time-locked cortical activity, thereby enabling investigation of oscillatory dynamics under controlled conditions. TMS can induce a brief increase in the power of oscillatory activity that is thought to share a common neurophysiological origin with spontaneous endogenous cortical oscillations [9–11]. When applied over the motor cortex, TMS elicits transient synchronization of cortical neuronal activity within the beta band, among others [12,13]. Combining TMS with electroencephalography (EEG) thus offers to investigate beta band dynamics, non-invasively in humans, in a controlled way. However, recording TMS-evoked EEG responses is limited by the spatial resolution of the technique, contamination by non-brain sources and variability across subjects [14–16].

Corticospinal neurons are synchronized with ongoing cortical oscillations, which are conveyed as common oscillatory inputs to lower motor neurons (MN) [17–20]. When MN are recruited, these common inputs can be transmitted in an approximately linear manner by the MN pool, enabling muscle activity to act as a peripheral readout of cortical oscillations [21– 24]. TMS combined with EEG can emulate this process, and probe connections between the motor cortex and MNs innervating skeletal muscles [25]. Accordingly, electromyographic (EMG) recordings from muscles innervated by stimulated cortical regions may provide a complementary and physiologically grounded readout of TMS-induced corticospinal activity. Supporting this view, previous studies have shown that EMG can capture corticospinal oscillatory responses when cortical areas are indirectly engaged by external stimuli [26,27].

In this study, we use TMS-EEG and EMG to characterize oscillatory muscle responses evoked by focal TMS of the motor cortex, to assess the cortical origin of the induced muscle responses and whether the induced beta responses are related to endogenous beta rhythms. Moreover, by manipulating stimulation parameters, we assess whether the TMS-induced oscillatory EMG responses depend on the recruitment of subthreshold inhibitory circuits or on the engagement of excitatory inputs driving corticospinal output [28–31], characterize the transmission of TMS-evoked beta activity to a pool of MN.

## Materials and Methods

This study comprises three experiments. In Exp1 (N=18), force, EMG and EEG were combined to characterize cortical and muscular oscillatory responses to TMS and their dependence on stimulation parameters (Fig. 1A). In Exp2 (N=1), high-density surface and intramuscular EMG were used to examine how TMS-evoked oscillations project to a pool of MN (Fig. 1B). In Exp3 (N= 10), a cutaneous stimulation protocol was conducted to characterize muscle oscillatory responses following peripheral afferent stimulation (Fig. 1C) [32].

**Figure 1.**
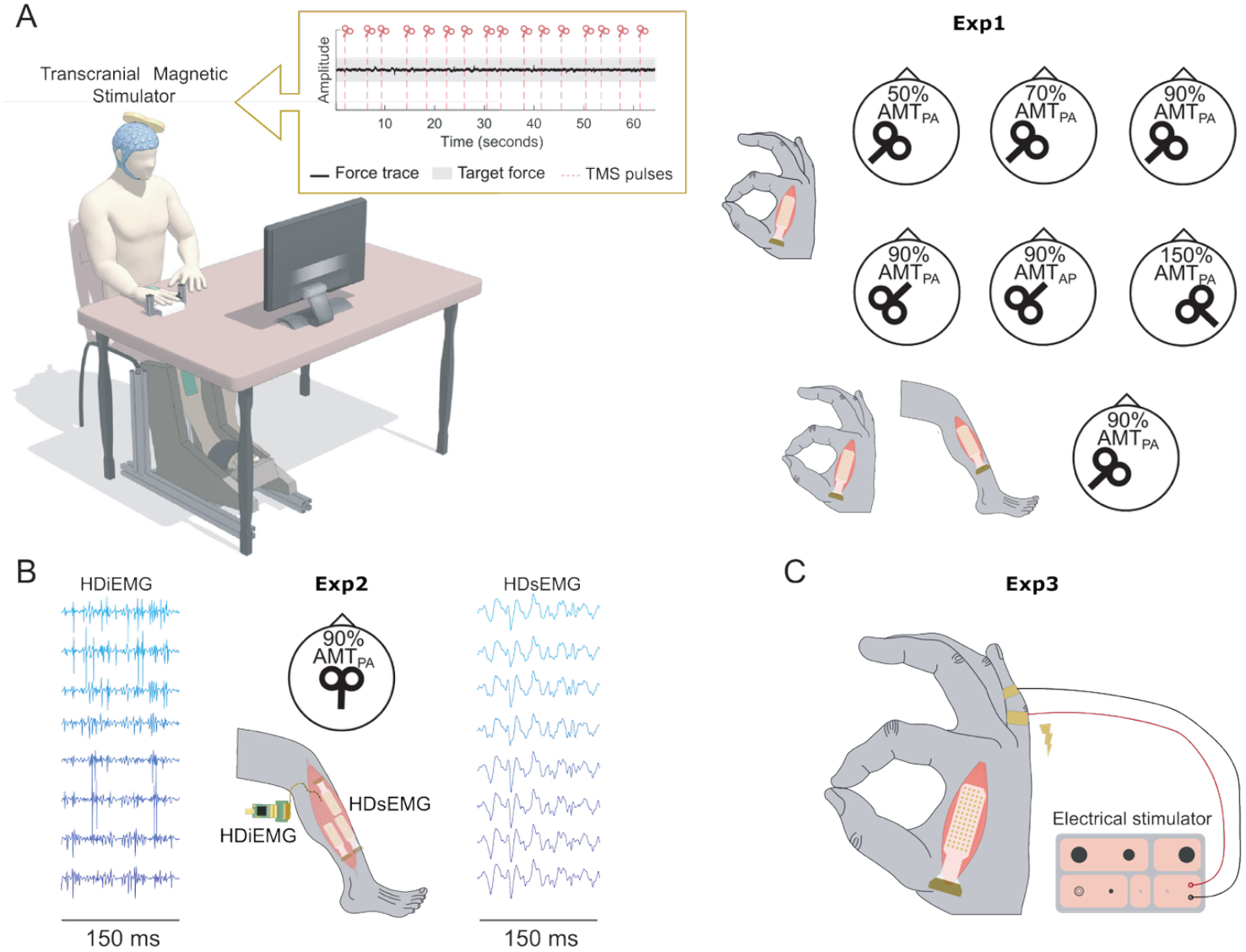
Experimental setups. **(A)** In Exp1, participants performed isometric contractions of the right first dorsal interosseous (FDI) muscle at 5% maximum voluntary contraction (MVC), either alone or combined with simultaneous right tibialis anterior (TA) contraction at 10% MVC. High-density surface EMG was recorded from both muscles. Each block consisted of multiple transcranial magnetic stimulation (TMS) pulses targeting the right FDI motor hotspot, except in one condition where the left FDI hotspot was targeted. TMS was delivered at various subthreshold intensities using either posterior-anterior (PA) or anterior-posterior (AP) coil orientations (summarized in the right panel). **(B)** In Exp2, TMS was applied over the right TA motor hotspot in the PA direction while participants maintained a contraction with the right TA at 10% of the MVC. High-density surface and intramuscular EMG were recorded simultaneously. The panel shows representative 150 ms segments from surface and intramuscular recordings. **(C)** Exp3 served as a control condition in which cutaneous electrical stimulation was delivered to elicit brief inhibition of FDI activity. This control demonstrated that peripheral stimuli produced comparable motor inhibition to what was obtained with TMS did not generate the oscillatory responses observed in the other experiments, supporting the cortical origin of the TMS-evoked responses.

The following sections describe the overall methods used (the detailed version of the methods is in the Appendix).

### Participants

All participants were healthy adults (N=28; 9 female; 20-40 years). Written informed consent was obtained prior to participation. All experiments were approved by the corresponding local ethics committees and conducted in accordance with the Declaration of Helsinki.

### Recordings

EEG was recorded from 63 scalp electrodes (10–20 system), referenced to FCz, grounded at FPz and sampled at 5kHz.

Force and high-density EMG were acquired with a Quattrocento amplifier during low-level isometric contractions: Exp1 involved the index finger abduction (EMG was obtained from the first dorsal interosseous -FDI-) and ankle dorsiflexion (EMG obtained from the right tibialis anterior -TA-); Exp2 involved ankle dorsiflexions (EMG of the TA); Exp3 involved a sustained pinching task (EMG of the FDI) (Fig. 1A,C). MVC was measured at session start and used to normalize force. The used force levels during the tasks were 5% MVC for FDI and 10% MVC for TA [40]. Surface EMG was recorded using 64-channel grids, band-pass filtered (20–500Hz) and sampled at 2042.48Hz.

In Exp2, in addition to surface EMG recordings using 3 64-channel grids, intramuscular EMG from TA was acquired with a 16-contact linear array (Fraunhofer Institute, Germany) at 10kHz [33].

All recording systems were synchronized via common digital triggers.

### Transcranial Magnetic Stimulation (TMS)

Single-pulse monophasic TMS was used. In Exp1, a figure-of-eight coil targeted the right FDI with posterior–anterior (PA) and anterior–posterior (AP) orientations. A condition with contralateral stimulation was included (Fig. 1A). In Exp2, a double-cone coil targeted the right TA with PA orientation. The hotspot was defined as the cortical location evoking the largest and most consistent Motor Evoked Potential (MEP) of the target muscle. Intensities were set relative to the active motor threshold (AMT) estimated with each stimulation condition.

### Cutaneous stimulation

In Exp3, peripheral nerve stimulation was achieved using an isolated constant current stimulator (DS3, Digitimer Ltd. UK) through two stainless steel ring electrodes placed around the first and the second phalanges of the fifth digit. Pulse width was 200μs and intensity was set to 10× each participant’s sensory threshold [32].

### Experimental Design

Exp1 and Exp2 investigated oscillatory activity elicited by TMS in muscles (Fig. 1A-B). First, MVC was assessed for each recorded muscle. Subsequently, the positions of the TMS coil and the AMT for the FDI (Exp1) or TA (Exp2) were determined. During the recording blocks, subjects maintained isometric contractions. TMS pulses were delivered on average every 5s with a 20% temporal jitter.

Exp1 started with eight blocks involving 40 TMS pulses delivered over the right FDI hotspot in the PA direction. The first two blocks were at 90% AMT with both FDI and TA contracted. The next six blocks were with intensities at 50%, 70%, and 90% AMT (two blocks per intensity) with only the FDI contracted. Two additional blocks of 60 pulses in the AP direction were performed either at 90% AMT estimated using AP stimulation (effect-matched) or at 90% AMT estimated using PA stimulation (intensity-matched). Finally, a block of 60 pulses was delivered over the left FDI hotspot at 150% AMT estimated for the right FDI using PA orientation.

In Exp2, three blocks of 30 TMS pulses were delivered over the right TA hotspot in the PA direction while the participant performed an isometric contraction of the TA.

In Exp3, cutaneous electrical stimuli were delivered to the fifth digit at intervals of 1.8±0.2s while participants maintained a sustained isometric contraction of the right FDI at 10% MVC (Fig. 1C) [32].

### Data analysis

Force signals were normalized to MVC and resampled to 2048Hz. Data were band-pass filtered (0.5–100Hz) and notch filtered at 50Hz. Signals were segmented into 2s epochs around the stimulus, and noisy trials were excluded.

In Exp1, EMG was resampled to 2048Hz and notch filtered at 50Hz. Bad channels were excluded, and data were segmented into 2s epochs around the TMS pulse. Trials containing MEPs or excessive noise were excluded. Remaining signals were rectified and averaged across channels.

EEG was processed using EEGLAB and TESA toolbox [34,35]. Continuous recordings were merged, noisy channels excluded, and data segmented into 2 s epochs around the TMS trigger with baseline correction. TMS-related artefacts were initially reduced by excising and interpolating the stimulus period, followed by downsampling and trial rejection. Two rounds of independent component analysis (fastICA [36,37]) removed residual artefacts. Finally, signals were band-pass filtered (1–100Hz), notch filtered (48–52Hz), and re-referenced to a common average.

Time–frequency analyses were performed using the FieldTrip toolbox, with pre-processed data detrended and decomposed using a multitaper approach.

Spectral corticomuscular coherence was computed after applying a surface Laplacian to EEG [21] and detrending both signals. Short-time Fourier transforms were calculated trial-by-trial. Cross- and auto-spectral estimates were then pooled across trials to derive complex coherence. For each participant, the EEG channel around C1 with highest coherence was used for time–frequency coherence maps and statistics [21].

In Exp2, EMG was decomposed into MN spike trains using a blind source separation (Swarm-Contrastive Decomposition) [45,46,47]. Prior to decomposition, EMG was downsampled to 2kHz, band-pass filtered (10–500Hz), and line noise removed (49–51Hz). Intramuscular EMG was band-pass filtered (150–4500Hz). 60 s segment from one block was selected for decomposition, retaining MNs with silhouette values >0.88 [40]. Identified MN were tracked across the recording, manually reviewed [41], and units active in fewer than 30 trials were excluded. Instantaneous firing rates were computed from MN spike trains. Neural drive to the muscle was estimated using the cumulative spike train (CST) [42]. Peristimulus time histograms (PSTHs) were computed from MN spikes using 1-ms bins aligned to TMS [43].

To assess TMS-evoked activity in the pool of MNs, time-frequency analysis was applied to the CST. Before that, the CST was band-pass filtered in the beta band (13-35Hz [44]) and averaged across TMS-stimuli, following the same pipeline described above. The resulting spectrum was used to identify the time and frequency of the maximal response (*t*_*ß*_, *f*_*ß*_). Beta projections onto the MN pool were characterized by repeating this analysis on CSTs built from increasing numbers of randomly selected MNs. Peak power was extracted for each pool size (mean of 50 random iterations).

TMS-evoked responses were then analysed at the level of individual MNs. Spikes from each MN were summed across trials (CST_MN_), band-pass filtered f_ß_ (±2Hz) and the instantaneous amplitude computed via Hilbert transform. Response power change was quantified as the RMS of a 100 ms window centred on *t*_*ß*_, expressed relative to the RMS of 1 s pre-stimulus period. To compare MNs of different sizes (“onion skin” principle [45]), each CST_MN_ was normalized by selecting by selecting trials to equalize total spike counts [46]. We finally assessed the relationship between firing rate and power change.

In Exp3, occasional stimulation artifacts in FDI EMG (segment of ±10ms around the simulus) were replaced with Gaussian noise based on pre-stimulus amplitude, followed by 20– 500Hz band-pass filtering. Time–frequency analysis was then performed using the same procedure as in Exp1.

### Statistics

Significant changes in EEG and EMG (time and time–frequency domains) were assessed using a cluster-based permutation approach in FieldTrip [47], with a cluster-defining threshold of p < 0.05 and a minimum of two channels for EEG. Permutations were performed across channels and time windows of interest, and corrected two-tailed p<0.05 was considered significant. Analyses typically compared the interval [0,0.6] s post-stimulus with the interval [−1,−0.1] s which was considered the baseline within each condition.

To examine the relationship between endogenous beta activity and TMS-evoked responses, corticomuscular coherence was calculated before (averaging four consecutive pre-stimulus windows of 200ms between −800ms and 0ms) and after TMS (the first 200ms). The EEG channel with maximal beta coherence per subject was used. Normality was checked and Pearson correlations were computed, with one-tailed p<0.05 considered significant.

Beta power differences in individual MNs between pre- and post-TMS windows were assessed with paired t-tests (after Shapiro-Wilk normality check, two-tailed p<0.05). The relationship between MN firing rate and beta power change was evaluated with a linear regression, reporting R^2^ and model p-value.

## Results

Overall, on average, 21±8% of the TMS trials were discarded per subject due to bad EMG or EEG trials (see SuppTable 1). The percentage of trials in which an MEP was observed across conditions and subjects was 9±10%.

The estimated AMTs in the PA and AP directions were 45±5% and 57±11% of the maximum stimulator output.

### TMS induces a salient beta response in tonically active muscles

To assess the characteristics of the TMS-induced cortical oscillations transmitted to the muscles, we first analysed the recordings from Exp1 involving PA TMS targeting the right FDI while the right FDI and TA were contracted. Fig. 2 shows the EEG and EMG responses. EEG responses were in line with previous publications. In the time domain, the well-documented P30, N45, P60, N100 and P200 TMS-evoked potentials could be observed around the stimulated brain area (Fig. 2A) [9,48]. A significant cluster of increased power was found in the 2-26Hz range and 15-285ms interval (*p*=0.015) (Fig. 2B). A later cluster was found in the interval 454-600ms and in the 2-7Hz range (*p*=0.015), likely related to oscillatory responses to salient sensory stimuli [49].

**Figure 2.**
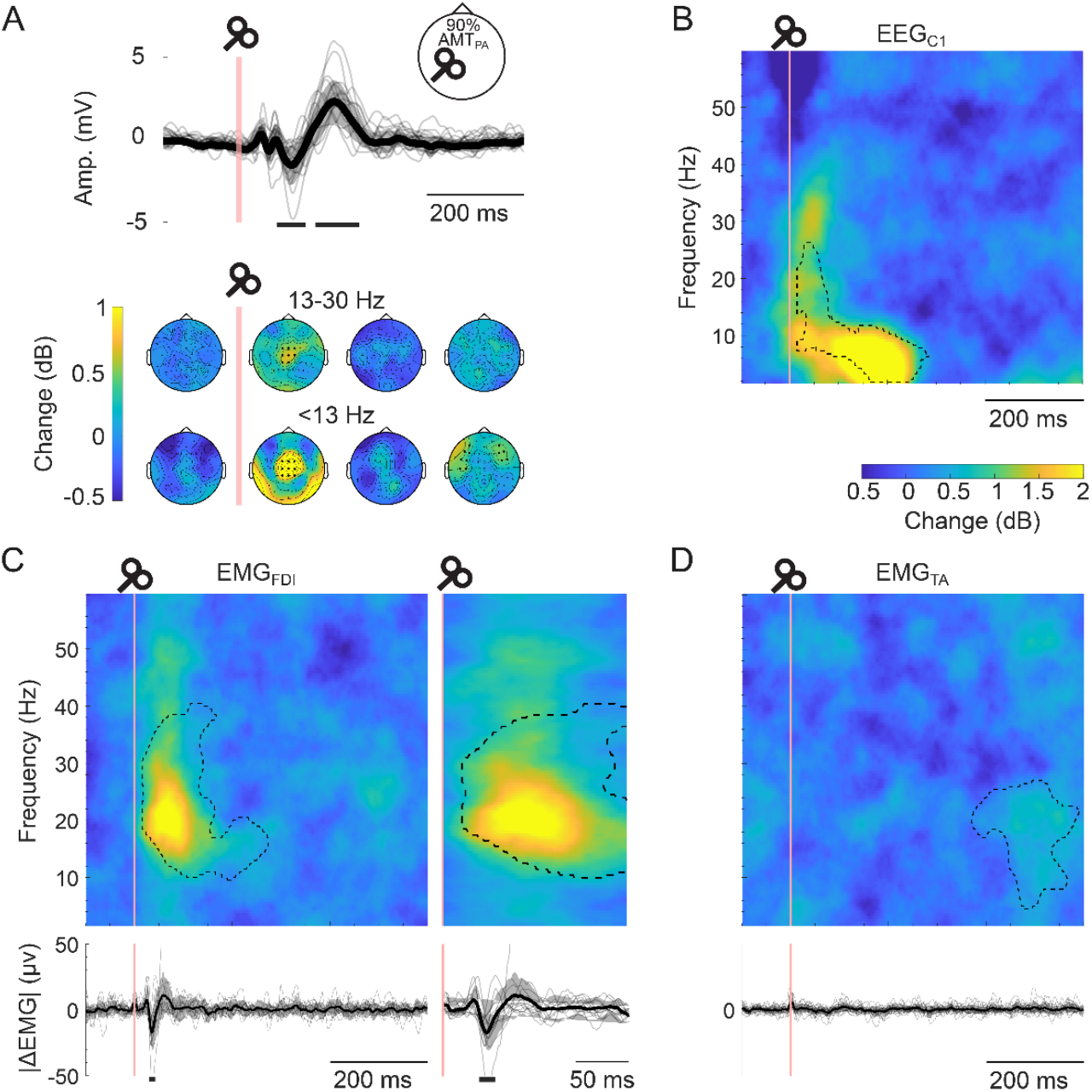
EEG and EMG responses from the right FDI and TA to sub-motor threshold TMS stimulation over the right FDI hotspot at 90% of the AMT in the PA direction (first condition from Exp1). **(A)** TMS-evoked potentials measured with EEG. The top panel shows the evoked EEG activity in C1. The scalp maps show power changes across 200 ms segments before and after TMS; **(B)** Time-frequency map of the average EEG activity in C1; **(C)** Time-frequency map (left) of the average HDsEMG signal from the FDI (top) and changes in amplitude relative to pre-TMS intervals of the rectified EMG (bottom); time-frequency map and rectified EMG on the right represents a zoomed window around the TMS-evoked beta response; **(D)** Same as C but for the TA. Significant clusters in the time-frequency maps are represented with dashed lines. Significant deviations from the baseline level in the time plots are marked with horizontal black lines beneath the plots. Also in the time domain plots, grey lines represent individual participants and black lines represent the average across participants. Scale of power change (dB) is similar across B, C and D. Significance threshold was set at p < 0.05.

In the EMG, no clear changes were observed below 10Hz, but a significant cluster in the beta band was identified within 9-40Hz in the interval 15-274ms after the TMS (*p*=0.007) (Fig. 2C). The cluster showed a peak at 21Hz and 65ms. In terms of frequency, this is in line with the typically observed frequency of beta oscillations transmitted between motor cortex and muscles [21]. Considering that the expected corticomuscular beta transmission latency for FDI is approximately 21ms a cortically generated beta response in FDI would be expected to have its peak power over the motor cortex roughly 40-50 ms after the stimulus onset. Indeed, Fig. 2B reflects an increase around 20Hz matching this temporal range.

In the time domain, the rectified EMG from the FDI showed a marked decrease peaking at 35ms followed by an increase above pre-TMS levels peaking around 60ms (Fig. 2C bottom). This is in line with previous studies assessing the effects of sub-motor threshold TMS on active muscles [50]. These early inhibitory and excitatory EMG responses may contribute to the initial part of the observed beta increase in the time-frequency maps, although they alone cannot explain the observed maximum at later latencies.

The EMG from the TA muscle did not show oscillatory responses early after (0-150 ms) the TMS stimulus (Fig. 2D). The observed brief inhibition-facilitation sequence in the rectified EMG in the FDI was not observed in the TA either (Fig. 2D bottom). However, the TA did present a significant cluster of increased power between 4-27Hz and 370-594ms and peaking at 17Hz (p=0.002). This later beta response is in line with previously described increases in corticospinal beta power ∼500ms after salient sensory stimuli [27].

To investigate a possible cortical origin of the beta response in the FDI, we calculated the corticomuscular coherence (CMC) between EEG and EMG (Fig. 3A). The temporal evolution of coherence showed an increase in coherence in the 0-200ms interval in the cortical areas beneath the coil. A significant cluster showing an increase in coherence between 13-30Hz and 0-200ms was found involving channels around the stimulated area (Fig. 3A-B). To test for a relationship between endogenously generated cortical beta transmitted to muscles during sustained contractions and TMS-evoked beta, we assessed whether CMC measured during the contraction periods preceding the TMS pulses was correlated with the CMC post-TMS (Fig. 3C). A significant correlation (*p*=0.003) was observed between these measures, indicating that the participant-dependent level of beta CMC post-TMS was linearly related to CMC levels obtained during sustained contractions without stimulation. This suggests that TMS-evoked beta responses share the same cortical generators as the ones involved in the generation of endogenous beta activity in the motor cortex.

**Figure 3.**
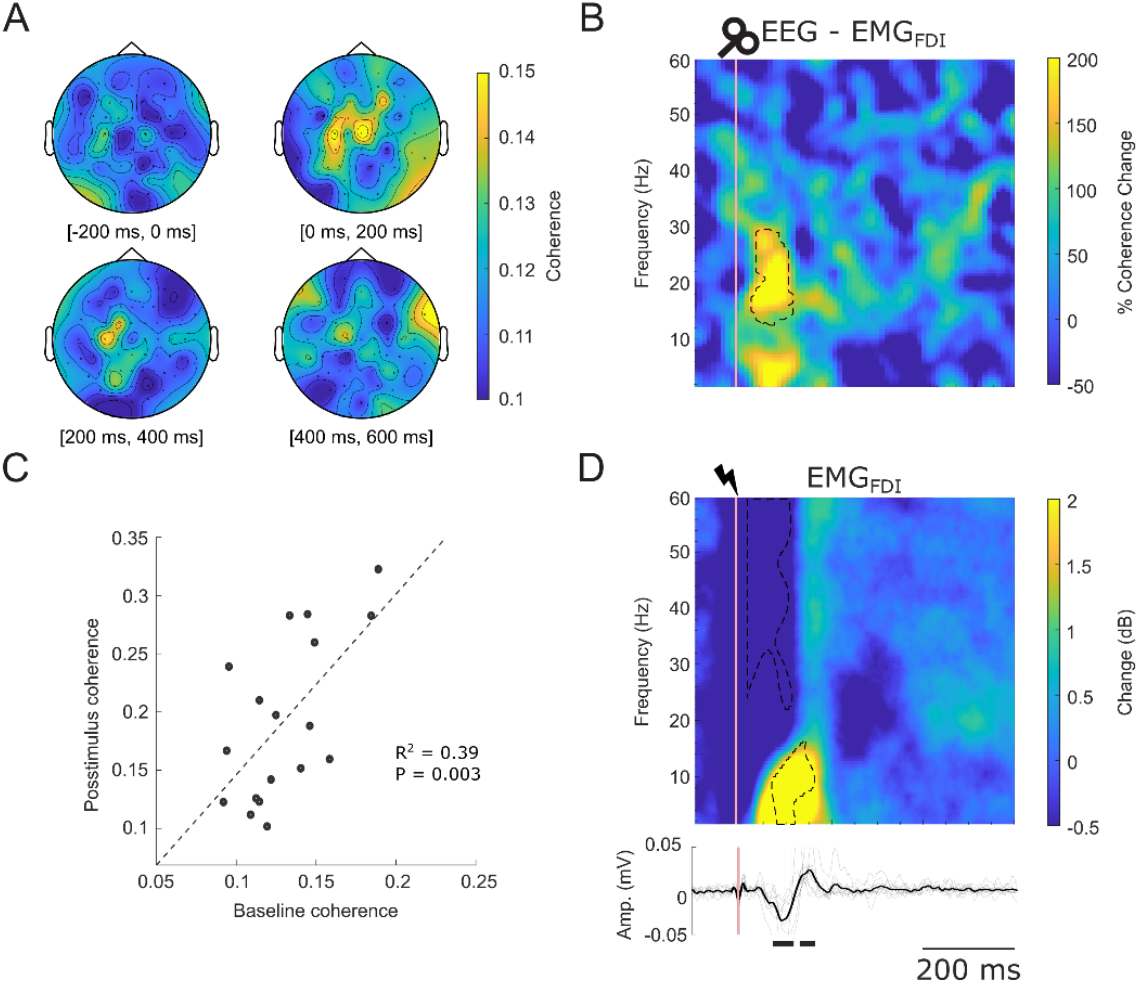
**(A)** Scalp map of corticomuscular coherence with the FDI across four non-overlapping windows of 200 ms; **(B)** Time-frequency representation of corticomuscular coherence changes after TMS (choosing the optimal EEG channel for each subject). A significant cluster of coherence increase is observed between 33-125ms and 12-29Hz; **(C)** Correlation between each participant’s beta coherence amplitude before the TMS (computed as the average of three non-overlapping 200 ms windows) and in the 0-200 ms window after TMS; **(D)** Time-frequency map of changes in EMG power after cutaneous stimuli are delivered producing a brief inhibitory input to the MN pool (Exp3). In panels B and D, the pink vertical lines represent the time at which stimuli were delivered. Dashed lines in panels B and D represent the limits of the identified clusters were significant changes in power are observed. Significance threshold was set at p < 0.05.

Finally, we also included a control experiment (Exp3) to assess the time-frequency response in the FDI when cutaneous stimulation is delivered causing a brief inhibition of the MNs [32]. Fig. 3D shows the spectral responses obtained. The results show a prominent increase at low frequencies (1-16Hz) that is related to the induced inhibition. The obtained significant cluster (*p=*0.010) starts 69ms after the electrical stimuli and has a duration of 93ms [32]. On the contrary, there were no observable increases at frequencies around the beta band. A cluster of reduced power at high frequencies soon after the stimulus was also observed, related to the induced silencing of MNs.

### TMS-evoked beta responses in the EMG depend on stimulus intensity and activated intracortical circuits with the TMS

To investigate how different cortical inputs driving corticospinal output influenced the induction of cortical beta activity transmitted to muscles, we performed experiments changing the intensity, orientation and location of the TMS.

First, we compared EMG responses to stimuli delivered at 50%, 70% and 90% of the AMT (Fig. 4 A-C). When 90% was used, a significant cluster of enhanced beta activity post-TMS was observed (15-316ms, 1-60Hz, *p*=0.002), as seen in the previous section. In addition, significant intervals of inhibition (32-42ms; *p*=0.005) and facilitation (57-75ms; *p*=0.005) were observed in the rectified EMG. On the contrary, 50% and 70% conditions did not result in visible changes in the time-frequency maps right after stimulation (although 50% condition resulted in a mild but significant increase around 258-453ms and 14-31Hz; *p*=0.020). None of the two conditions showed a significant change in the rectified EMG post-TMS.

**Figure 4.**
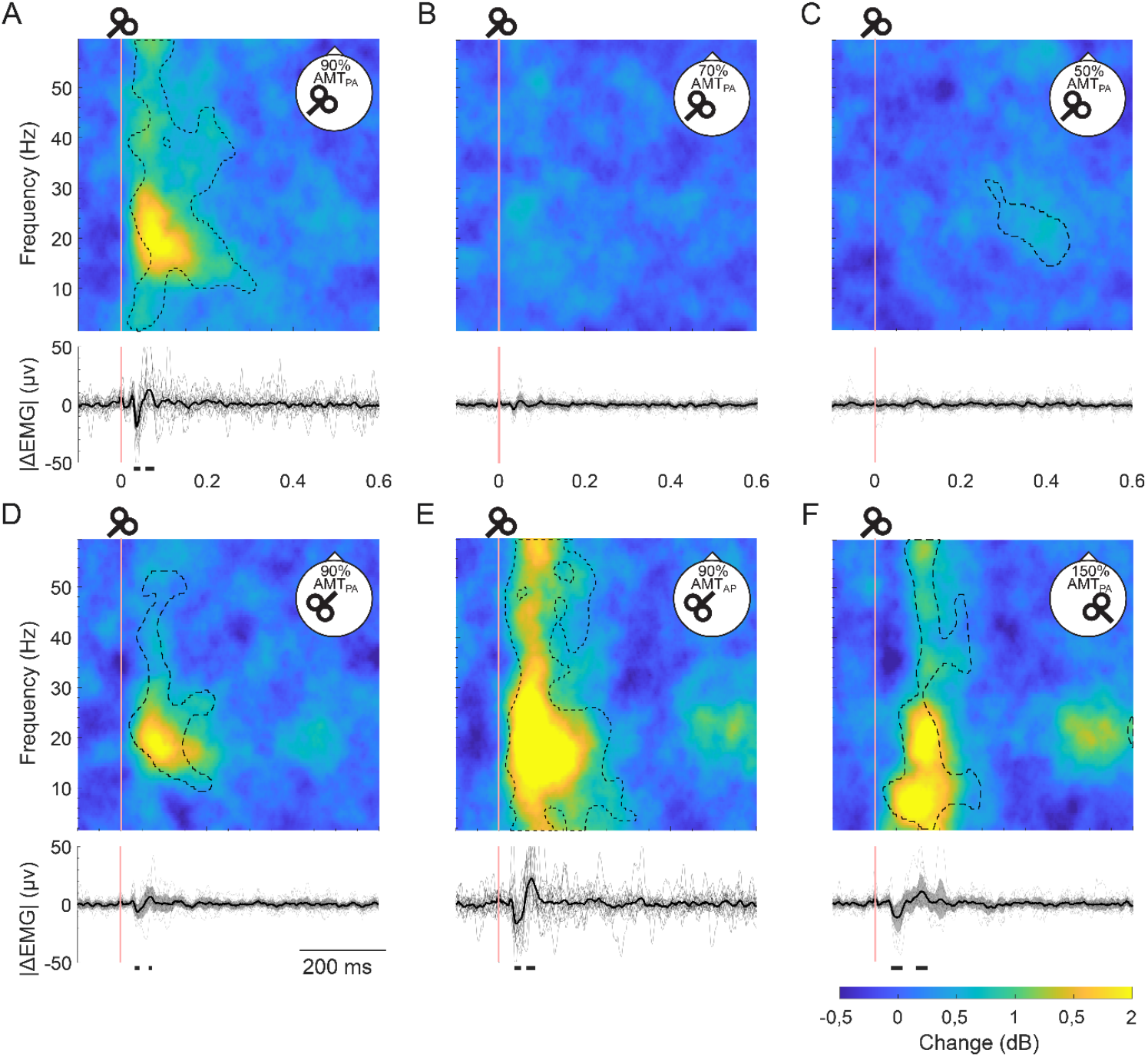
Right FDI responses to magnetic stimulation. For each condition tested, the panel shows a time-frequency map (top) and the changes in the rectified EMG signal relative to pre-TMS intervals (bottom). **(A-C)** Reults obtained using intensities at 90%, 70% and 50% of the AMT and stimuli delivered over the left motor cortical representation of the right FDI using PA orientation. **(D-E)** Reults obtained delivering TMS pulses in the AP direction over the right motor cortex with an intensity at either 90% of the AMT in the PA direction (D) or 90% of the AMT in the AP direction (E). **(F)** TMS delivered at 150% of the AMT in the PA direction with the coil targeting the motor cortical hotspot for the left FDI in the PA direction. Significant clusters in the time-frequecy maps are marked using dashed lines. In the time plots of the rectified EMG, significant changes post-TMS are marked with solid horizontal black lines beneath the x axes. The time plots at the bottom of the panels represent both participant-specific results (grey lines) and avarge results (black lines). Significance threshold was set at p < 0.05.

Then we conducted blocks of stimulation with the TMS coil oriented in the AP direction using either intensity-matched stimulation (same intensity as with PA direction) or effect-matched stimulation (using 90% of the AMT with AP direction). In the intensity-matched case, stimulation resulted in a significant cluster in the beta band peaking at 81ms and 19Hz (cluster extension: 20-219ms, 9-53Hz, *p*=0.005) (Fig. 4D). The rectified EMG revealed a significant inhibitory effect at 35–43ms (*p=*0.010), followed by a significant facilitation at 67–73ms (*p=*0.020). With effect-matched stimulation, a cluster reflecting a strong and broad beta response was obtained peaking at 17Hz (cluster extension: 15-322ms, 1-60Hz, *p*=0.002) (Fig. 4E). The rectified EMG showed significant inhibition (36-51ms, *p*=0.005) followed by facilitation (66-83ms, *p*=0.002).

Finally, the contralateral stimulation condition led to a significant cluster in the 17-244Hz range (showing two different peaks at 6Hz and 88ms and at 20Hz and 118ms; *p*=0.005). The average rectified EMG showed a significant inhibition peaking at 46ms (*p=*0.005) followed by a significant facilitation peaking at 104ms (*p*=0.005). A significant cluster at later latencies was also found, likely related to the saliency of the stimulus used (502-800ms, 9-27Hz, *p*=0.017).

### Sampling of TMS-evoked cortical beta oscillations by motor neurons

A total of 34 MNs were decomposed and tracked across blocks in Exp2 (mean discharge rate 12.7±1.9 spikes/s). Due to subtle variations in the force generated and in the activity of the MNs, each MN could be characterized in a different number of trials. On average, we were able to track the individual MN in 88±13 trials per MN (range 64-98).

Fig. 5A shows the TMS-evoked activity in the PSTH. A brief inhibitory period was observed with a latency of 33ms, representing a global decrease in the firing occurrence. Fig. 5B shows the spike trains from all trials of all the MNs segmented around the TMS and the CST after band-pass filtering (13-35 Hz) and averaging across trials. The resulting signal showed an oscillation at 20Hz peaking at 105ms and with a duration of approximately 100ms (Fig. 5C).

**Figure 5.**
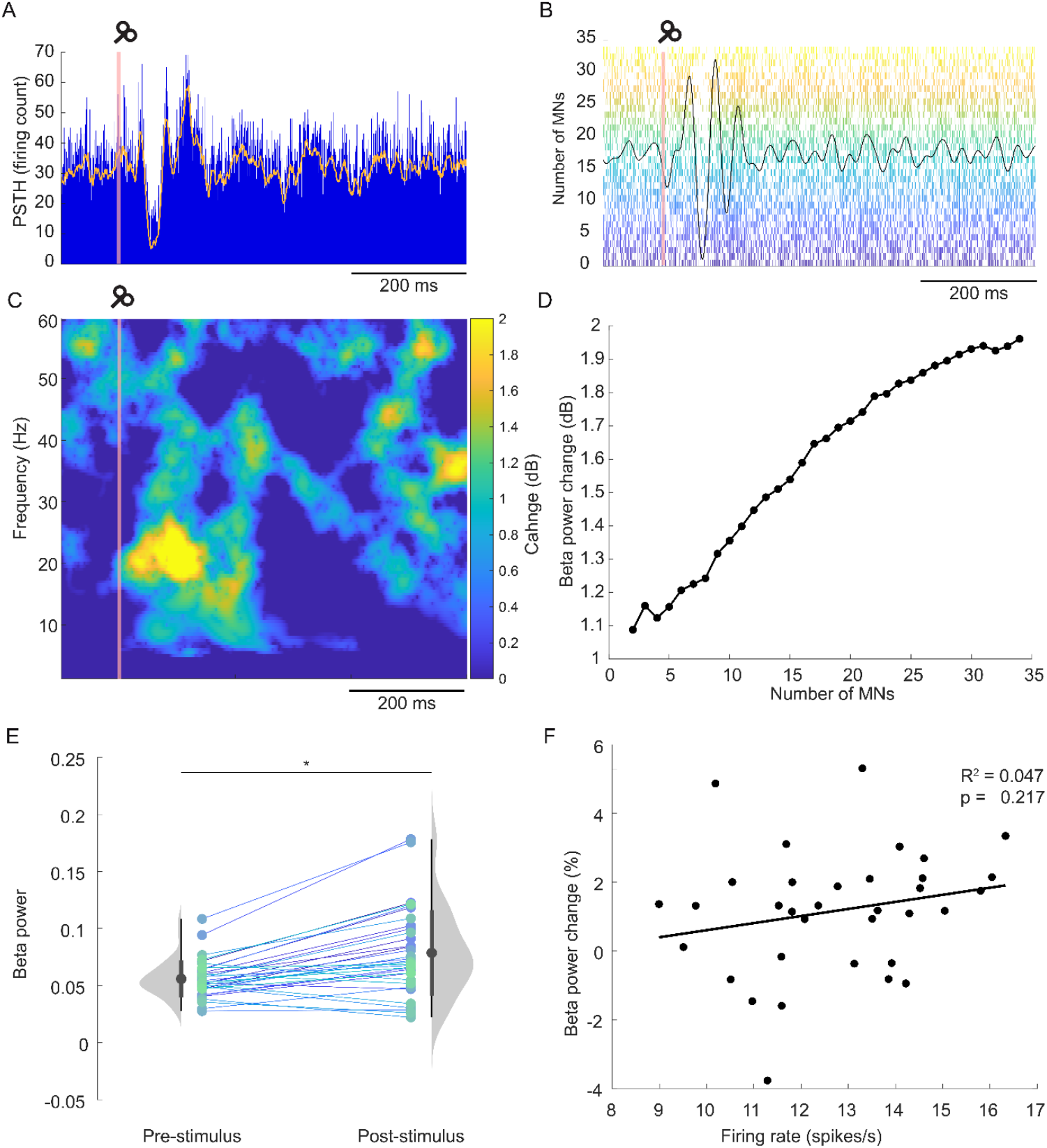
**(A)** Peri-stimulus time histogram of all MUs detected and tracked before and after the TMS stimuli. The orange line represents the filtered version of the histogram; **(B)** Band-pass filtered version of the composite spike train obtained by summing all the MUs decomposed and steadily firing. The panel also shows the firings of each motor unit in the back (each row shows all the spikes in all the trials in which a motor unit has been detected and tracked); **(C)** Time-frequency analysis of power changes in the composite spike train (CST) obtained by summing the 34 MUs decomposed. The largest post-stimuli power change peaks at 105ms and 20 Hz. (**D**) Post-stimuli power change in the beta band as a function of the number of MUs included in the analysis. (**E**) Instantaneous power in the beta band for individual MUs (individual-coloured dots) in pre- and post-stimulus windows. (**F**) Post-stimuli instantaneous power change (%) in the beta band for individual MUs and their mean firing rate. Significance threshold was set at p < 0.05 (*).

Then we repeated the time-frequency analysis on subsets of increasing numbers of randomly selected MN. The analysis revealed that beta power increased as a function of the number of MN considered (Fig. 5D). The linear rate of increase of this curve indicates that the TMS-evoked response was likely projected as a common input to the MN pool [22].

We also estimated the power of the evoked beta response in individual MN within a 100-ms window centred at 105ms and filtered at 20±2Hz (the time and frequency of the peak beta power). Compared to pre-stimulus levels, individual MN significantly increased their beta power post-TMS (0.056±0.016 vs 0.079±0.037, *p<*0.001, Fig. 5E). No relationship was found between TMS-evoked beta power and MN firing rate (R^2^=0.047, *p=*0.217) (Fig. 5F).

## Discussion

This study shows that subthreshold TMS over the primary motor cortex evokes corticospinal beta-band oscillations that are related to endogenous cortical beta activity and can be measured from muscles. These findings establish a novel framework to probe corticospinal beta oscillations with precision and circuit specificity.

TMS over motor cortex evoked oscillatory activity in several frequency bands at the cortical level, but selectively induced a beta response entraining corticospinal neurons. Our study is aligned with previous primate research showing that stimulation of pyramidal tract neurons evokes beta oscillations in cortical local field potentials and in activated muscles controlled by the stimulated cortical regions [18]. In both cases, the origin of the beta response is likely related to the engagement of inhibitory interneuron circuits projecting onto corticospinal neurons, which induce an early inhibition of motor outputs reflected in the reduction of the EMG amplitude [50]. This post-stimulus inhibition represents the starting point of the beta response in both studies and is consistently observed here in all TMS conditions that lead to EMG beta responses.

Manipulation of stimulation intensity and coil orientation reveals how these inhibitory circuits are accessed. With PA stimulation, beta oscillations appeared only near AMT, suggesting that a certain amount of stimulation is required to recruit the inhibitory interneuronal networks responsible for beta generation. In contrast, AP stimulation elicited beta oscillations at the same absolute intensity as PA stimulation, despite this lying well below AMT and being likely insufficient for direct corticospinal activation [30]. This demonstrates that beta generation reflects engagement of inhibitory circuits with lower thresholds and reduced directional sensitivity compared to excitatory corticospinal inputs [51]., when intensity was adjusted relative to motor threshold in each direction, AP stimulation produced temporally and spectrally broader beta responses.

These findings indicate that stimulation direction excitatory corticospinal recruitment, while beta generation reflects recruitment of direction-insensitive inhibitory circuits [51]. Practically, AP stimulation may be advantageous for measuring corticospinal beta oscillations, as it engages inhibitory circuits while minimizing direct corticospinal activation. Contralateral stimulation further supports the involvement of common inhibitory circuits, eliciting beta responses with similar temporal and spectral characteristics to direct stimulation. TMS delivered over the contralateral hemisphere is known to engage transcallosal inhibitory pathways, and here it elicited beta responses with temporal and spectral characteristics similar to those observed with direct stimulation. The approximately 40ms delay in beta peak latency suggests involvement of transcallosal pathways and additional intracortical inhibitory processing [52].

Several findings in our study support that TMS-induced beta oscillations relate to endogenous cortical beta activity, providing a controlled approach for studying corticospinal interactions. First, TMS-induced oscillations increased corticomuscular beta coherence, with individual coherence amplitudes correlating with pre-TMS beta coherence during sustained contractions. This suggests individuals with stronger endogenous corticospinal beta coupling exhibit stronger TMS-evoked responses, consistent with a shared cortical generator. Second, MN analyses revealed that TMS-evoked beta oscillations are transmitted as uniform common input across the active MN pool, mirroring the projection pattern of endogenous beta activity during voluntary contractions [46]. Together, these findings indicate that TMS transiently entrains physiological corticospinal circuits rather than generating an artificial oscillatory response.

Overall, the obtained results motivate a broader reflection on the functional role of beta activity in the motor system. In the present experimental context, beta oscillations appear to emerge as an automatic, reflex-like response to a brief perturbation of motor cortical activity. This raises the possibility that beta oscillations may act as a stabilizing or resetting mechanism, reinstating a default or baseline state of motor cortical activity following transient disruptions [8]. Under this view, beta activity may represent a reactive process that restores network stability following transient disruptions. Such a mechanism could also operate under natural conditions, where spontaneous or externally driven transient events disrupt intermittently ongoing cortical dynamics [7,53].

At the same time, the present results indicate that beta activity occupies a privileged position in corticospinal communication. Although TMS evokes oscillatory activity across a broad frequency range at the cortical level, only beta oscillations are consistently transmitted to the muscles. Lower-frequency oscillations, such as those around 10 Hz, were not propagated to the motor periphery despite being present cortically. This selective transmission implies the existence of frequency-specific filtering mechanisms within the corticospinal system.

Several limitations should be acknowledged. All experiments were performed during sustained isometric contractions, a context known to facilitate the transmission of corticospinal beta oscillations. Consequently, the conclusions drawn here are specific to active and steady motor states. In addition, the MN decomposition experiment was conducted in a single participant due to the technical complexity, invasiveness, and cost. Nevertheless, the ability to reliably track a large population of MN provides a robust basis for the conclusions drawn.

## Conclusions

Subthreshold TMS over motor cortex induces transient corticospinal beta oscillations that can be reliably detected in muscle activity during sustained contractions. These responses share key features with endogenous corticomuscular beta rhythms and are broadly transmitted across the active motor neuron pool, supporting a common cortical origin. EMG recordings thus provide a sensitive and physiologically grounded approach to probe TMS-evoked brain oscillations and their corticospinal transmission.

