## Appendix for "TMS-Evoked Corticospinal Beta Oscillations in Humans Recorded from Muscles"

#### Extended Materials and Methods

This study comprises one main experiment and two additional experiments. In the main experiment (referred to as Exp1; sample size N=18 participants) we combined surface EMG in with EEG to characterize the transmission of TMS-induced oscillatory cortical activity to muscles (Fig. 1A). In this experiment, we also assessed the effects of different stimulus configurations, including variations in stimulus intensity, coil orientation, and stimulation site, on the induced responses. In a second experiment (Exp2; N=1), we combined surface and intramuscular EMG using high-density multielectrode arrays, enabling the decomposition of individual activity of a large number of motor neurons (MNs, Fig. 1B). This approach allowed us to characterize how TMS-induced cortical oscillatory activity is projected to individual MNs. Finally, we reanalysed data from a previous study (Exp3; N=10) to characterize muscle oscillatory responses elicited by peripheral nerve electrical stimulation (Fig. 1C) [34]. This experiment provided a control condition to assess whether oscillatory EMG activity observed after TMS was also present following peripheral afferent stimulation. This comparison was particularly relevant because both stimulation modalities induce a brief inhibition of MN activity [35–37], which could potentially influence the estimation of oscillatory responses in muscle recordings.

##### Participants

Eighteen healthy individuals were recruited for Exp1 (6 female, 3 left-handed, age: 26±4). One participant from Exp1 also took part in Exp2 (male, age: 41). None of the participants had a clinical history of neurological, musculoskeletal, or psychiatric disorders, and none were taking medications affecting the central nervous system. The two experiments were approved by the local ethics committee (Comité de Ética de la Investigación de la Comunidad Autónoma de Aragón: CEICA; IDs: PI23_404 and PI23-546).

Data from Exp3 were obtained from ten healthy participants (3 female, age: 30±5). The study was approved by Imperial College London ethics committee (reference number 18IC4685).

All participants recruited for this work provided written informed consent prior to the experimental procedures. All the experiments were conducted in accordance with the Declaration of Helsinki.

##### Force measurements

In Exp1, forces exerted by the right FDI and right TA were measured. Participants rested their right arm on a table with the hand pronated and the index finger positioned against a force cell (FC22, Measurement Specialties, USA) measuring index finger abduction force generated by FDI contraction (see Fig. 1A). Ankle dorsiflexion forces were recorded using a custom-made platform designed to isolate ankle movement, with force measured by a CCT TF022 transducer and amplified using a Forza-b device (OT Bioelettronica, Torino, Italy). Participants received continuous visual feedback of the force produced by both muscles.

In Exp2, ankle dorsiflexion forces were recorded using the same configuration as in Exp1.

In Exp3, participants performed a pinching task between the index finger and the thumb of the right hand (Fig. 1C). Forces applied by the index finger were measured with a load cell (FC22, Measurement Specialties, US).

In all experiments, the maximum voluntary contraction (MVC) of the investigated muscles was measured at the beginning of the experimental sessions. MVC was estimated from three maximal isometric contractions, each lasting approximately 3–5 seconds, and calculated as the average of the peak force values obtained across the three trials for the index finger or ankle. Force signals were digitized together with EMG recordings using a Quattrocento device sampling at 2042.48Hz (OT Bioelettronica, Torino, Italy).

**Electroencephalography (EEG)**

Electroencephalography (EEG) was acquired in Exp1 to characterize the TMS-evoked cortical responses and to relate them to the responses observed in the recorded muscles. A DC-coupled, TMS-compatible EEG system was used (ActiChamp, Brain Products GmbH, Germany). The EEG signal was acquired from 63 active electrodes embedded in an elastic cap (actiCAP) and arranged according to the international 10–20 system. FCz and FPz positions served as reference and ground electrodes, respectively. Electrode-skin impedances were maintained below 10kΩ for all channels and the sampling frequency was 5kHz throughout the recording session.

**Electromyography (EMG)**

In Exp1, high-density surface EMG was obtained from the FDI and TA. Grids consisting of 64 electrodes were placed over the FDI and TA muscles of the right hand and foot. For the FDI, a grid of 5×13 electrodes with 4mm inter-electrode distance was used, whereas for the TA, a 5×13 grid with 8mm inter-electrode distance was used. Two wet wristbands were used as reference and grounding electrodes on the right wrist for the FDI and on the right ankle for the TA (grounds were short-circuited).

In Exp2, a thin-film intramuscular high-density electrode with 16 recording contacts was inserted in the TA muscle. The electrode consisted of a polyimide substrate carrying a linear array of platinum recording electrodes (5,257$\mu m^2$ per contact) with an inter-electrode distance of 1 mm (Fraunhofer Institute, Germany). Electrode insertion was guided by ultrasonography and performed using a 25G guiding needle with the electrode embedded inside. Once placed within the target portion of the muscle, the needle was withdrawn, leaving the electrode inside the muscle belly [38]. Additionally, three EMG grids (5×13 electrodes) with 4 mm inter-electrode distance were placed over the TA muscle, covering as much of the muscle surface as possible while maintaining a distance of approximately 2 cm from the intramuscular insertion site.

In Exp3, surface EMG was recorded from the right FDI using a 5 × 13 electrode grid with an inter-electrode distance of 4 mm. Two wet wristbands placed on the right wrist served as reference and ground electrodes.

Surface EMG signals were recorded in monopolar configuration, band-pass filtered (20-500 Hz) and sampled at 2042.48Hz (Quattrocento, OT Bioelettronica, Italy). Intramuscular EMG was recorded in monopolar mode at a sampling rate of 10 kHz using a multichannel amplifier (Open Ephys FPGA Acquisition Board, Lisbon, Portugal).

In all experiments, skin preparation before electrode placement included shaving when necessary, gentle abrasion with abrasive paste, and cleansing with 70% ethyl alcohol. Data acquired from the different recording systems were synchronized using common digital trigger signals sent to all the devices.

**Transcranial Magnetic Stimulation (TMS)**

Single-pulse, monophasic TMS was applied using a DuoMAG MP-Dual system (DEYMED Diagnostic s.r.o., Czech Republic). The device was connected to a 70mm figure-of-eight coil (70BF model) in Exp1 and to a 120mm double-cone coil (120BVFT model) in Exp2.

The hotspot for TMS was defined as the cortical location within the primary motor cortex where stimulation evoked the largest and most stable motor evoked potential (MEP) in the target muscle.

In Exp1, the target muscle was the right FDI. This muscle was stimulated either via its contralateral cortical representation or via stimulation of the ipsilateral motor cortex, in which case coil placement was guided by MEPs recorded from the left FDI. Accordingly, three stimulation hotspots were identified: (1) the hotspot for the right FDI with the coil oriented in the postero-anterior (PA) direction (*i.e.*, with the handle pointing backwards and leftwards forming an angle of approximately 45º with the midline); (2) the hotspot for the right FDI with the coil oriented in the antero-posterior (AP) direction (*i.e.*, with the coil rotated 180º relative to the PA condition); and (3) the hotspot for the left FDI using the coil in the PA orientation (*i.e.*, same as in the first case but with the handle pointing rightwards) [31,32,39].

In Exp2, the target muscle was the right TA, and the stimulation hotspot was determined using a PA coil orientation.

In all experiments, stimulation intensities were set relative to the active motor threshold (AMT) of the target muscle. The AMT was defined as the minimum stimulation intensity required to elicit a visible MEP (typically exceeding ~200µV peak-to-peak amplitude, depending on baseline EMG activity) in 5 out of 10 consecutive stimuli trials during a tonic isometric contraction. Contraction levels were 5% MVC for the FDI and 10% MVC for the TA. This difference was based on the distinct motor unit properties of the two muscles and were chosen to recruit sufficiently large motor neuron pools while minimizing fatigue [40].

**Cutaneous stimulation**

In Exp3, peripheral nerve stimulation was achieved using an isolated constant current stimulator (DS3, Digitimer Ltd. UK) through two stainless steel ring electrodes placed around the first and the second phalanges of the fifth digit. Stimulation pulse width was set to 200 µs, and conductive gel was applied between the ring electrodes and the skin to reduce impedance. The sensory threshold, defined as the minimum stimulation intensity that generated a subjective perception on the finger, was estimated for each participant. The stimulus intensity during the actual recordings was set to 10x the sensory threshold [34].

**Experimental Design**

Exp1 investigated oscillatory activity elicited by TMS in muscles (Fig. 1A). Participants completed a single session comprising multiple experimental conditions. At the beginning of the session, MVC was assessed for each recorded muscle. Subsequently, the positions of the TMS coil and the AMT of the right FDI (Exp1) or right TA (Exp2) were determined. During the recording blocks, subjects maintained an isometric contraction of the FDI (5% of the MVC) and, in some cases, also the TA (10% of the MVC). Single TMS pulses were delivered on average every 5 seconds with a 20% temporal jitter to reduce predictability.

The first two recording blocks involved the delivery of 40 TMS pulses in the PA direction over the right FDI hotspot at 90% of the AMT, while the right FDI and TA were contracted. These recordings allowed us to assess and compare EEG and EMG oscillatory activity post-TMS and to compare responses between the FDI (targeted by the TMS) and the TA (non-targeted).

Subsequent blocks were performed while only the FDI was contracted. To study the influence of the TMS intensity on the induced EMG responses, TMS was delivered in the PA direction over the right FDI hotspot at three intensities: 50%, 70%, and 90% AMT. Each intensity was tested in pairs of blocks involving 40 TMS pulses each.

To investigate the contribution of different interneuronal circuits to the induced EMG responses, additional blocks were conducted using an AP TMS direction [31–33]. One block was delivered at 90% of the AMT estimated using AP stimulation, and a second block at 90% of the AMT estimated using PA stimulation, allowing comparison of intensity-matched and effect-matched AP and PA conditions. This design allowed us to test whether TMS-induced activity depends primarily on the engagement of corticospinal output, in which case EMG responses would be expected when AP stimulation reaches its own corticospinal threshold, or instead on the recruitment of specific directionally sensitive cortical inputs, in which EMG oscillatory responses would be expected at lower absolute stimulation intensities or selectively for PA stimulation. For these conditions, a single block of 60 pulses was recorded to limit total experiment duration.

Finally, a block of 60 TMS pulses was delivered over the left FDI hotspot at 150% of the AMT estimated in the right FDI using PA orientation. This condition was included to assess oscillatory EMG responses elicited by activation of motor cortical regions projecting to the right FDI via transcallosal pathways.

In Exp2, we investigated how TMS-induced corticospinal oscillations are transmitted to populations of individual MNs within an active pool. TMS was delivered over the cortical representation of the right TA, with EMG recorded from the same muscle. The participant maintained isometric ankle dorsiflexion at 10% MVC. Three recording blocks were performed, each consisting of 30 TMS pulses delivered in the PA direction at 90% AMT, with the same inter-pulse interval as in Exp1.

Finally, in Exp3 we assessed EMG oscillatory responses elicited by peripheral stimulation. Participants maintained a sustained precision pinch between the thumb and index finger at 10% MVC during a 200-s recording period, while EMG was recorded from the FDI. Cutaneous electrical stimuli were delivered to the fifth digit at intervals of 1.8±0.2s. This protocol elicited the cutaneous silent period, a well-characterized withdrawal reflex mediated by A-delta afferents projecting to spinal interneurons and producing transient inhibition of motor neurons innervating intrinsic hand muscles (Fig. 1C) [35].

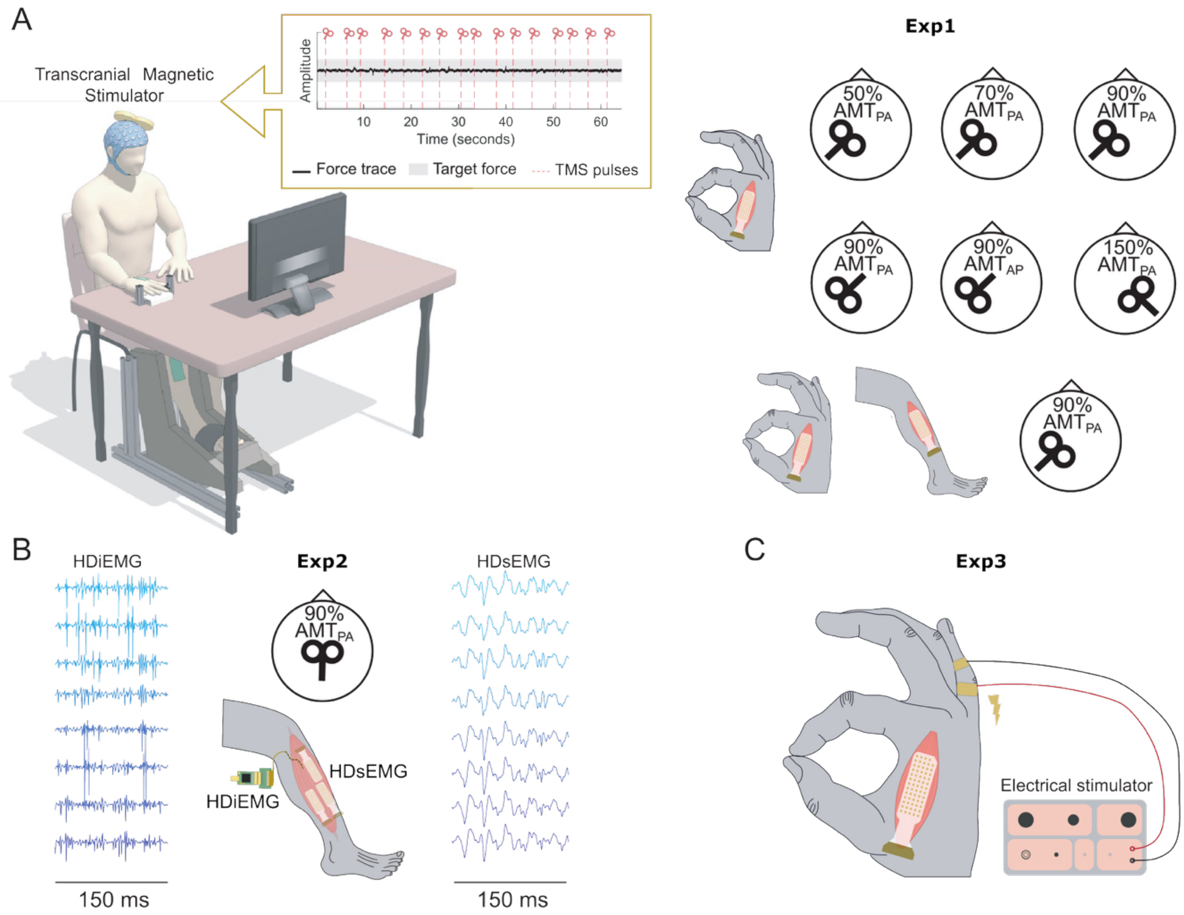


***Figure 1.-*** *Experimental setups.* ***(A)*** *In Exp1, participants performed isometric contractions of the right first dorsal interosseous (FDI) muscle at 5% maximum voluntary contraction (MVC), either alone or combined with simultaneous right tibialis anterior (TA) contraction at 10% MVC. High-density surface EMG was recorded from both muscles. Each block consisted of multiple transcranial magnetic stimulation (TMS) pulses targeting the right FDI motor hotspot, except in one condition where the left FDI hotspot was targeted. TMS was delivered at various subthreshold intensities using either posterior-anterior (PA) or anterior-posterior (AP) coil orientations (summarized in the right panel).* ***(B)*** *In Exp2, TMS was applied over the right TA motor hotspot in the PA direction while participants maintained a contraction with the right TA at 10% of the MVC. High-density surface and intramuscular EMG were recorded simultaneously. The panel shows representative 150 ms segments from surface and intramuscular recordings.* ***(C)*** *Exp3 served as a control condition in which cutaneous electrical stimulation was delivered to elicit brief inhibition of FDI activity. This control demonstrated that peripheral stimuli produced comparable motor inhibition to what was obtained with TMS did not generate the oscillatory responses observed in the other experiments, supporting the cortical origin of the TMS-evoked responses.*

**Data analysis**

All analyses were conducted offline within the MATLAB environment (Version 2024a; MathWorks Inc., Natick, USA).

Force signals were normalized within each participant based on the prior MVC estimation. This transformation mapped the reference contraction level (e.g., 5% MVC) to zero, and all reported force variations correspond to deviations relative to this baseline.

Signals were resampled to 2048Hz before further processing. Data were filtered between 0.5 and 100Hz using a fourth-order zero-phase Butterworth filter, and a 50Hz notch filter was applied to supress line noise. Signals were then segmented into 2s epochs (from -1s to 1s with respect to the TMS pulses). Trials in which the maximum variation of the smoothed force signal exceeded the mean across trials plus three standard deviations were excluded from subsequent analysis.

In Exp1, high-density surface EMG signals were resampled to 2048Hz. Line noise at 50Hz was attenuated using a second-order infinite impulse response (IIR) notch filter centered at 50Hz (quality factor Q=35), applied in a zero-phase manner using forward–backward filtering. Bad channels were identified as those with an amplitude standard deviation exceeding the mean standard deviation across all channels by more than three standard deviations and were excluded from subsequent analyses. These channels were removed prior to analysis. EMG trials were extracted from the continuous recordings and segmented from −1s to 1s relative to TMS pulses. To determine whether an MEP was present in the trial, we used a custom-designed script which first compared the average of all trials with a standard reference MEP in terms of latency and waveform morphology, thereby defining a subject-specific MEP template. Each individual trial was then compared against this template and discarded if the characteristic latency and waveform were visually detected. Trials were considered excessively noisy when they exhibited abrupt or non-physiological amplitude fluctuations. Finally, the EMG signals were rectified and averaged across channels for further analysis.

EEG analysis was conducted using EEGLAB in combination with functions from the TMS–EEG Signal Analyser (TESA) MATLAB toolbox [41,42]. Continuous EEG data from multiple runs were first merged. Noisy channels, defined by the presence of excessive artefacts or abnormal signal fluctuations, were identified by visual inspection and excluded from further analysis. A channel was considered noisy if it showed excessive artefacts or abnormal signal fluctuations. The data were then segmented into epochs from −1s to 1s relative to the TMS trigger, and each epoch was baseline-corrected by subtracting its mean value. A standard procedure was applied to remove TMS-related artefacts. First, the initial TMS artefact was removed by excising EEG signals from −5ms to 10ms around the stimulus and interpolating the removed segment using a cubic function from −1ms to 1ms over the gap. Signals were then downsampled, and bad trials, defined as those with pronounced artefacts, were removed following visual inspection. Artefact removal was further refined using independent component analysis (ICA) with the fastICA algorithm [43,44]. At this stage, only components associated with large-amplitude, early artefacts (including TMS-evoked muscle and voltage decay) were discarded. Subsequently, data removal was extended to −5ms to 15ms, with cubic interpolation applied over −5ms to 5ms around the gap. The data were then filtered using a band-pass filter (1–100 Hz) and a notch filter (48–52Hz), both fourth-order Butterworth. Remaining artefacts (mainly eye movements, blinks or EMG artifacts) were eliminated through a second ICA round and component selection, resulting in clean EEG signals. Finally, the cleaned signals were re-referenced to a common average reference for subsequent analysis.

Time–frequency analyses were performed using the FieldTrip toolbox for MATLAB (version 20240417). Pre-processed data were detrended using a polynomial removal procedure. Time–frequency decomposition was carried out with the multitaper convolution method, employing a Hanning taper and a sliding window of 200ms with a 10ms step size.

Spectral coherence was computed from the auto-spectral and cross-spectral estimates of EEG and EMG signals. EEG signals were first transformed using a surface

Laplacian (for each recording position, the average amplitude of the nearest channels was subtracted [24]) and both EEG and EMG were linearly detrended prior to spectral estimation. Then, for each EEG channel and for the EMG, the short-time Fourier transforms were obtained on a trial-by-trial basis using sliding windows of 200ms (10ms steps) (MATLAB's *spectrogram* function was used with a spectral resolution of 0.5Hz). This yielded complex-valued short-time spectra $S_x(f,t,i,j)$ for EEG (with $f$, $t$, $i$ and $j$ denoting the frequency, time, channel and trial dimensions) and $S_y(f,t,j)$ for EMG. Then, for each trial and EEG channel, cross-spectral and auto-spectral estimates were derived as:

$$S_{xy}(f,t) = S_x(f,t,i,j)\, S_y^*(f,t); S_{xx}(f,t) = |\, S_x(f,t,i,j)\, |^2; S_{yy}(f,t) = |\, S_y(f,t)\, |^2$$

These quantities were computed for each time–frequency bin and stored across trials and channels. Coherence was then obtained by pooling spectral estimates across trials (*i.e.*, summing trial-wise cross- and auto-spectra) and normalizing:

$$C_{xy}(f,t,i) = \frac{\sum_{\mathrm{j}} S_{xy}\,(f,t,j,i)}{\sqrt{\left(\sum_{\mathrm{j}} S_{xx}\,(f,t,i,j)\right)\left(\sum_{\mathrm{j}} S_{yy}\,(f,t,j)\right)}}$$

This procedure yields a complex coherence estimate whose magnitude reflects the strength of coupling between EEG channels and the EMG at each frequency and time window. The channel yielding the highest coherence magnitude among those around the stimulated area (C3, C1, Cz, FC1, CP1) for each participant was used to compute the average time-frequency maps of corticomuscular coherence and to run statistical tests aimed at finding significant changes in coherence across time and frequencies [24].

In Exp2, EMG was automatically decomposed into MN spike trains using a blind source separation technique [45,46]. Specifically, we applied Swarm-Contrastive Decomposition (SCD), a method designed to isolate the activity of individual MNs by leveraging the spatiotemporal characteristics of motor unit action potentials and the inherent sparsity of MN firing patterns [46,47]. Prior to decomposition, surface EMG was first downsampled to 2 kHz, band-pass filtered (10-500Hz, zero-phase 2$^{nd}$ order Butterworth) and line noise was removed with a band-stop filter (49-51Hz, 2$^{nd}$ order Butterworth). Intramuscular EMG was band-pass filtered (150-4500Hz, zero-phase 2$^{nd}$ order Butterworth). Due to the high computational cost of the automatic decomposition, a segment of 60s of one of the three TMS blocks was selected for decomposition. MNs with a silhouette value (SIL) above 0.88 were retained for the analysis. The SIL value is a normalized indicator of the robustness in the decomposition, and a value of 0.88 indicates a separation by approximately 2.75 standard deviations between spikes and background activity [45]. The resultant MNs were tracked across the rest of the recording by reapplying the separation filters obtained from the decomposition [45]. MN spike trains were then manually reviewed and edited correcting for errors derived from the automatic procedure [48]. MNs that were not active during at least 30 or more trials were discarded from further analysis.

From the spike trains, the instantaneous firing rate was computed for individual MNs. As an estimation of the neural drive transmitted to the muscle, the cumulative spike train (CST) was computed with the aggregated activity of individual MNs over time [49]. Peristimulus time histogram (PSTH) was constructed by binning the MN spike trains (1-ms bin width) and aligning them around the TMS-stimuli [50].

To explore the TMS-evoked oscillatory activity in the pool of MNs decomposed, the time-frequency analysis was applied over the CST. Before that, the CST was band-pass filtered in the beta band (13-35 Hz, zero-phase 2nd order Butterworth, [51]) and averaged across TMS-stimuli, following the same pipeline described above for non-decomposed EMG signals. The resulting time-frequency spectrum was used to determine the time and frequency at which a maximum response was obtained (referred to as $t_ß$ and $f_ß$).

To characterize the projection of beta oscillations to the MN pool, time-frequency analysis was iteratively performed on CSTs comprising an increasing number of randomly selected MNs. The peak power at ($t_ß, f_ß$) was obtained for each MN pool size (50 iterations with random MN selections were performed, and the average power was computed).

TMS-evoked responses were then analysed at the level of individual MNs. To do this, the spiking activity of each (i) MN was summed across trials resulting in a CST per MU ($CST_{MNi}$). The resulting signals were band-pass filtered at $f_ß$ (±2Hz, zero-phase 2nd order Butterworth). The instantaneous amplitude of the resulting signals was computed using the Hilbert transform, and the response power for each MN was estimated as the root mean square within a 1 s pre-stimulus baseline window and a 100ms window centred around $t_ß$.

Then, we investigated how TMS-evoked beta responses projected into MNs of different sizes (based on firing rate and recruitment threshold relationship according to the onion skin principle [52]). The sampling resolution of each $CST_{MNi}$ previously computed depends on the total number of spikes included, which is determined by the number of trials and the firing rate of the MNs. To ensure comparable sampling resolution across MNs with different firing rates, we normalized each MN $CST_{MN}$ by selecting the number of trials such that the total spike count was similar across MNs [53]. We finally assessed the relationship between the firing rate of individual MNs and the rate of power change between the baseline and the TMS-evoked window. This normalization prevented the analysis from being biased by differences in sampling resolution across MUs.

In Exp3, EMG from the FDI was ocassionally contaminated with the electrical stimulation artifact. Prior to analysis, a segment of ±10ms around the simulus was replaced with simulated gaussian noise with the mean amplitude of pre-stimulus EMG signal channel by channel, and then the signals were band-passed filtered (20-500 Hz, zero-phase $2^{nd}$ order Butterworth). After this preprocessing step, the same analysis procedure as in Exp1 was applied for the time-frequency analysis.

**Statistics**

To look for significant changes in the EEG and EMG signals (both in the time and time-frequency domains) statistical analyses were conducted using a permutation approach implemented in the open-source MATLAB toolbox FieldTrip [54]. Correction for multiple comparisons was applied with a cluster-based algorithm using t-tests as the test statistic. Clusters were ranked according to the sum of t-values across all points within each cluster. A cluster-defining threshold of $p < 0.05$ was used. A minimum cluster size of two channels was used for EEG time-frequency maps. Permutations were performed in the channel × time domain across all recorded channels and within the time windows of interest. Corrected two-tailed p-values < 0.05 were considered statistically significant. For most analyses, pairwise comparisons were carried out between the time interval of interest (ToI; 0 to 0.6 s relative to stimulus onset) and a baseline period (−1 to −0.1 s) within the same condition.

In order to study the relationship between endogenously generated beta activity and the TMS-induced beta responses, we analyzed the correlation between the corticomuscular coherence observed before and after the TMS stimuli. To obtain the pre-

and post-TMS coherence levels, the same methodology described above was applied, restricted to specific time windows. Pre-TMS coherence was obtained by averaging values across four consecutive pre-stimulus intervals [-800, -600] ms, [-600, -400] ms, [-400, -200] ms and [-200, 0] ms. Post-TMS coherence was estimated in the [0, 200] ms window. The EEG channel leading to larges corticomuscular coherence in the beta band in each subject was used for this analysis. Using the resulting mean values (pre- and post-), normality was assessed and a Pearson correlation was computed. One-tailed p-values < 0.05 were considered statistically significant.

Finally, to assess the power difference in the beta band for individual MNs between the pre-TMS window and the post-TMS window, we applied paired t-tests (normality of the data was priorly assessed with the Shapiro-Wilk test), considering statistical significance as two-tailed p-value < 0.05. To study the relationship between the MN firing rate and the power change in the beta band due to the TMS-evoked activity, a linear regression model was built using the MN firing rate as predictor and the power change as dependent variable. The goodness of fit of the linear relationship was estimated with $R^2$ and the model p-value.

#### Discarded trials

##### Supplementary table 1. Proportion (%) of Discarded Trials in Exp1

| | PA direction Right FDI hotspot | | | | | | AP direction Right FDI hotspot | | PA direction Left FDI hotspot |
|---|---|---|---|---|---|---|---|---|---|
| | FDI + TA | | | FDI | | | FDI | | FDI |
| Subj | 90% AMT$_{PA}$ | | | 50% AMT$_{PA}$ | 70% AMT$_{PA}$ | 90% AMT$_{PA}$ | 90% AMT$_{PA}$ | 90% AMT$_{AP}$ | 150% AMT$_{PA}$ |
| | EMG | EEG | CCM | EMG | EMG | EMG | EMG | EMG | EMG |
| 1 | 21% | | | 0% | 0% | 6% | 6% | 17% | 0% |
| 2 | 25% | 3% | 25% | 0% | 0% | 15% | 15% | 27% | 0% |
| 3 | 24% | 4% | 20% | 0% | 0% | 15% | 10% | 13% | 0% |
| 4 | 18% | 1% | 21% | 0% | 0% | 8% | 14% | 17% | 0% |
| 5 | 20% | 0% | 14% | 0% | 0% | 19% | 8% | 10% | 0% |
| 6 | 14% | 1% | 10% | 0% | 0% | 15% | 17% | 15% | 0% |
| 7 | 9% | 1% | 25% | 0% | 0% | 8% | 8% | 8% | 0% |
| 8 | 15% | 0% | 31% | 0% | 0% | 11% | 15% | 18% | 0% |
| 9 | 12% | 13% | 32% | 0% | 0% | 6% | 0% | 13% | 0% |
| 10 | 15% | 3% | 20% | 0% | 0% | 11% | 7% | 23% | 0% |
| 11 | 15% | 1% | 10% | 0% | 0% | 11% | 3% | 13% | 0% |
| 12 | 20% | 0% | 15% | 0% | 0% | 24% | 23% | 33% | 0% |
| 13 | 25% | 3% | 18% | 0% | 0% | 10% | 8% | 18% | 0% |
| 14 | 6% | 5% | 25% | 0% | 0% | 13% | 7% | 7% | 0% |
| 15 | 18% | 1% | 26% | 0% | 0% | 28% | 2% | 22% | 0% |
| 16 | 31% | 0% | 6% | 0% | 0% | 42% | 8% | 40% | 0% |
| 17 | 19% | 4% | 23% | 0% | 0% | 19% | 17% | 28% | 0% |
| 18 | 30% | 10% | 36% | 0% | 0% | 31% | 3% | 25% | 0% |
| Mean | 19% | 3% | 21% | 0% | 0% | 16% | 10% | 19% | 0% |
| SEM | 7% | 4% | 8% | 0% | 0% | 10% | 6% | 9% | 0% |

#### AMTs and TMS intensities

##### Supplementary table 2. Active Motor Thresholds (AMTs) and Stimulation Intensities in Exp1

| | AMT | | PA direction Right FDI hotspot | | | | AP direction Right FDI hotspot | | PA direction Left FDI hotspot |
|---|---|---|---|---|---|---|---|---|---|
| | | | FDI + TA | FDI | | | FDI | | FDI |
| Subj | PA | AP | 90% $AMT_{PA}$ | 50% $AMT_{PA}$ | 70% $AMT_{PA}$ | 90% $AMT_{PA}$ | 90% $AMT_{PA}$ | 90% $AMT_{AP}$ | 150% $AMT_{PA}$ |
| 1 | 44 | 56 | 40 | 22 | 31 | 40 | 40 | 51 | 66 |
| 2 | 50 | 83 | 45 | 25 | 35 | 45 | 45 | 75 | 75 |
| 3 | 44 | 59 | 40 | 22 | 31 | 40 | 40 | 53 | 66 |
| 4 | 38 | 51 | 34 | 19 | 27 | 34 | 34 | 46 | 57 |
| 5 | 42 | 46 | 38 | 21 | 29 | 38 | 38 | 41 | 63 |
| 6 | 40 | 57 | 36 | 20 | 28 | 36 | 36 | 51 | 60 |
| 7 | 45 | 48 | 41 | 23 | 32 | 41 | 41 | 43 | 68 |
| 8 | 48 | 63 | 43 | 24 | 37 | 43 | 43 | 57 | 72 |
| 9 | 40 | 77 | 36 | 20 | 28 | 36 | 36 | 69 | 60 |
| 10 | 45 | 48 | 41 | 23 | 32 | 41 | 41 | 43 | 68 |
| 11 | 42 | 55 | 38 | 21 | 29 | 38 | 38 | 50 | 63 |
| 12 | 45 | 49 | 41 | 23 | 32 | 41 | 41 | 44 | 68 |
| 13 | 36 | 46 | 32 | 18 | 25 | 32 | 32 | 41 | 54 |
| 14 | 47 | 60 | 42 | 24 | 33 | 42 | 42 | 54 | 71 |
| 15 | 44 | 47 | 40 | 22 | 31 | 40 | 40 | 42 | 66 |
| 16 | 53 | 60 | 48 | 27 | 37 | 48 | 48 | 54 | 80 |
| 17 | 49 | 53 | 44 | 25 | 34 | 44 | 44 | 48 | 74 |
| 18 | 54 | 76 | 49 | 27 | 38 | 49 | 49 | 68 | 81 |
| Mean | 45 | 57 | 40 | 23 | 32 | 40 | 40 | 52 | 67 |
| SEM | 5 | 11 | 4 | 3 | 4 | 4 | 4 | 10 | 7 |

### Statistical Summary Table

#### Supplementary table 3. Statistical Results for the Three Clusters with the Lowest p-values Identified in the EMG Time–Frequency Power Spectrum in Exp1

| | | | CLUS | SIGN | P-value | Range | | Peak | | |
|---|---|---|---|---|---|---|---|---|---|---|
| | | | | | | Time (ms) | Freq (Hz) | Time (ms) | Freq (Hz) | Change (dB) |
| PA direction Right FDI hotspot | 90% $AMT_{PA}$ FDI + TA | FDI | 1 | pos | **0.007** | 15-274 | 9-40 | 65 | 21 | 2.31 |
| | | | 2 | pos | 0.247 | 26-98 | 41-60 | 80 | 49 | 1.09 |
| | | | 3 | pos | 0.758 | 481-526 | 26-30 | 486 | 26 | 1.11 |
| | | TA | 1 | pos | **0.002** | 370-594 | 4-27 | 491 | 17 | 1.13 |
| | | | 2 | pos | 0.162 | 379-440 | 52-60 | 434 | 60 | 0.90 |
| | | | 3 | neg | 0.232 | 119-224 | 23-41 | 122 | 27 | 0.40 |
| | 50% $AMT_{PA}$ | FDI | 1 | pos | **0.020** | 258-453 | 14-31 | 402 | 21 | 1.02 |
| | | | 2 | neg | 0.323 | 529-598 | 37-48 | 557 | 45 | 0.40 |
| | | | 3 | pos | 0.667 | 296-327 | 45-50 | 321 | 47 | 0.62 |
| | 70% $AMT_{PA}$ | FDI | 1 | pos | 0.050 | 15-124 | 37-60 | 85 | 51 | 0.76 |
| | | | 2 | pos | 0.114 | 78-219 | 13-22 | 193 | 16 | 0.88 |
| | | | 3 | pos | 0.811 | 451-486 | 12-15 | 463 | 15 | 0.56 |
| | 90% $AMT_{PA}$ | FDI | 1 | pos | **0.002** | 15-316 | 1-60 | 76 | 19 | 2.15 |
| | | | 2 | pos | 0.479 | 448-514 | 17-25 | 461 | 17 | 0.91 |
| AP direction Right FDI hotspot | 90% $AMT_{PA}$ | FDI | 1 | pos | **0.005** | 20-219 | 9-53 | 81 | 19 | 2.22 |
| | | | 2 | neg | 0.448 | 253-307 | 36-45 | 260 | 36 | 0.52 |
| | | | 3 | pos | 0.781 | 205-221 | 5-7 | 205 | 6 | 0.97 |
| | 90% $AMT_{AP}$ | FDI | 1 | pos | **0.002** | 15-322 | 1-60 | 88 | 17 | 3.00 |
| | | | 2 | pos | 0.050 | 414-600 | 10-27 | 488 | 23 | 1.45 |
| | | | 3 | pos | 0.439 | 462-533 | 48-53 | 519 | 50 | 1.00 |
| PA direction Left FDI hotspot | 150% $AMT_{PA}$ | FDI | 1 | pos | **0.005** | 17-244 | 1-60 | 88 | 6 | 2.57 |
| | | | 2 | pos | **0.005** | 588-600 | 11-25 | 738 | 18 | 1.58 |
| | | | 3 | pos | 0.154 | 568-664 | 32-42 | 651 | 39 | 0.78 |

**Supplementary table 4. Statistical Results for the Three Clusters with the Lowest p-values Identified in the EEG Time–Frequency Power Spectrum in Exp1**

| | | CLUS | SIGN | P-value | Range | | Peak | | | |
|---|---|---|---|---|---|---|---|---|---|---|
| | | | | | Time (ms) | Freq (Hz) | Time (ms) | Freq (Hz) | Change (dB) | Chan |
| PA direction Right FDI hotspot | 90% $AMT_{PA}$ FDI + TA | 1 | pos | **0.015** | 15-285 | 2-26 | 194 | 2 | 3.96 | FCz |
| | | 2 | pos | **0.015** | 454-600 | 2-7 | 600 | 2 | 2.00 | AF7 |
| | | 3 | neg | 0.085 | 15-72 | 39-60 | 15 | 60 | -1.95 | CP6 |

**Supplementary table 5. Statistical Results for the Three Clusters with the Lowest p-values Identified in the Time–Frequency Representation of Corticomuscular Coherence in Exp1**

| | | CLUS | SIGN | P-value | Range | | Peak | | |
|---|---|---|---|---|---|---|---|---|---|
| | | | | | Time (ms) | Freq (Hz) | Time (ms) | Freq (Hz) | Change (dB) |
| PA direction Right FDI hotspot | 90% $AMT_{PA}$ FDI + TA | 1 | pos | **0.015** | 33-125 | 13-30 | 84 | 19 | 0.82 |
| | | 2 | pos | 0.436 | 43-115 | 2-7 | 74 | 4 | 0.63 |
| | | 3 | neg | 0.536 | 536-600 | 50-58 | 556 | 51 | -0.36 |

**Supplementary table 6. Statistical Results for the Three Clusters with the Lowest p-values Identified in the EMG Time–Frequency Power Spectrum in Exp3**

| | CLUS | SIGN | P-value | Range | | Peak | | |
|---|---|---|---|---|---|---|---|---|
| | | | | Time (ms) | Freq (Hz) | Time (ms) | Freq (Hz) | Change (dB) |
| Cutaneous Stimulation | 1 | pos | **0.010** | 69-162 | 1-16 | 112 | 6 | 5.26 |
| | 2 | neg | **0.010** | 15-114 | 22-60 | 15 | 24 | -1.79 |
| | 3 | pos | 0.701 | 422-450 | 33-36 | 434 | 36 | 0.37 |

**Supplementary table 7. Statistical Results for the Three Clusters with the Lowest p-values Identified in the EMG Time-Domain Analysis in Exp1**

| | | | CLUS | SIGN | P-value | Range Time (ms) | Peak Time (ms) | Peak Amplitude (µV) |
|---|---|---|---|---|---|---|---|---|
| PA direction Right FDI hotspot | 90% $AMT_{PA}$ FDI + TA | FDI | 1 | neg | **0.007** | 31-41 | 35 | -22.99 |
| | | | 2 | pos | 0.035 | 57-65 | 60 | 13.57 |
| | | | 3 | pos | 0.190 | 71-75 | 73 | 8.26 |
| | | TA | 1 | pos | 0.100 | 329-333 | 331 | 2.65 |
| | | | 2 | pos | 0.319 | 467-470 | 469 | 2.97 |
| | | | 3 | neg | 0.359 | 72-75 | 73 | -3.27 |
| | 50% $AMT_{PA}$ | FDI | 1 | pos | 0.214 | 101-103 | 103 | 4.68 |
| | | | 2 | pos | 0.383 | 472-474 | 473 | 3.72 |
| | | | 3 | pos | 0.527 | 94-96 | 96 | 4.84 |
| | 70% $AMT_{PA}$ | FDI | 1 | neg | 0.204 | 34-37 | 35 | -6.44 |
| | | | 2 | pos | 0.224 | 98-100 | 98 | 5.18 |
| | | | 3 | neg | 0.289 | 408-411 | 410 | -3.20 |
| | 90% $AMT_{PA}$ | FDI | 1 | pos | **0.005** | 57-75 | 63 | 14.27 |
| | | | 2 | neg | **0.005** | 32-42 | 35 | -24.70 |
| | | | 3 | pos | 0.047 | 24-29 | 26 | 14.60 |
| AP direction Right FDI hotspot | 90% $AMT_{PA}$ | FDI | 1 | neg | **0.010** | 35-43 | 40 | -8.42 |
| | | | 2 | pos | **0.020** | 67-73 | 72 | 8.82 |
| | | | 3 | neg | 0.463 | 163-166 | 164 | -3.70 |
| | 90% $AMT_{AP}$ | FDI | 1 | pos | **0.002** | 66-83 | 77 | 25.60 |
| | | | 2 | neg | **0.005** | 36-51 | 40 | -20.17 |
| | | | 3 | pos | 0.678 | 208-210 | 210 | 5.41 |
| PA direction Left FDI hotspot | 150% $AMT_{PA}$ | FDI | 1 | pos | **0.005** | 95-120 | 104 | 12.82 |
| | | | 2 | neg | **0.005** | 38-63 | 46 | -13.09 |
| | | | 3 | pos | 0.284 | 209-211 | 209 | 5.77 |

**Supplementary table 8. Statistical Results for the Three Clusters with the Lowest p-values Identified in the EEG Time-Domain Analysis in Exp1**

| | | CLUS | SIGN | P-value | Range Time (ms) | Peak Time (ms) | Peak Amplitude (mV) | Chan |
|---|---|---|---|---|---|---|---|---|
| PA direction Right FDI hotspot | 90% $AMT_{PA}$ FDI + TA | 1 | pos | **0.005** | 139-246 | 184 | 3.51 | Cz |
| | | 2 | neg | **0.005** | 61-234 | 94 | -2.70 | Cz |
| | | 3 | neg | **0.005** | 334-550 | 550 | -0.83 | AF8 |
